# Differential requirements for different subfamilies of the mammalian SWI/SNF chromatin remodeling enzymes in myoblast differentiation

**DOI:** 10.1101/2023.03.05.531193

**Authors:** Teresita Padilla-Benavides, Monserrat Olea-Flores, Tapan Sharma, Sabriya A. Syed, Hanna Witwicka, Miriam D. Zuñiga-Eulogio, Kexin Zhang, Napoleon Navarro-Tito, Anthony N. Imbalzano

## Abstract

Mammalian SWI/SNF (mSWI/SNF) complexes are ATP-dependent chromatin remodeling enzymes that are critical for normal cellular functions and that are mis-regulated in ∼20% of human cancers. These enzymes exhibit significant diversity in the composition of individual enzyme complexes. mSWI/SNF enzymes are classified into three general sub-families based on the presence or absence of specific subunit proteins. The three sub-families are called BAF (BRM or BRG1-associated factors), ncBAF (non-canonical BAF), and PBAF (Polybromo-associated BAF). The biological roles for the different subfamilies of mSWI/SNF enzymes are poorly described. We knocked down (KD) the expression of genes encoding subunit proteins unique to each of the three subfamilies, *Baf250A, Brd9*, and *Baf180*, which mark the BAF, ncBAF, and PBAF sub-families, respectively, and examined the requirement for each in myoblast differentiation. We found that BAF250A and the BAF complex was required to drive lineage-specific gene expression during myoblast differentiation. KD of *Baf250A* reduced the expression of the lineage determinant *Myogenin* and other differentiation markers, due to decreased binding of BAF250A to myogenic gene promoters. KD of *Brd9* delayed myoblast differentiation. However, RNA-seq analysis revealed that while the *Baf250A*-dependent gene expression profile included genes involved in myogenesis, the *Brd9*-dependent gene expression profile did not. Moreover, no-colocalization of Baf250A and Brd9 was observed in differentiating cells, suggesting independent mechanisms of action for BAF and ncBAF complexes in myogenesis. The PBAF complex was dispensable for myoblast differentiation. The results distinguish between the roles of the mSWI/SNF enzyme subfamilies during myoblast differentiation.

## Introduction

SWI/SNF chromatin remodelers are ATP-dependent enzymes that alter nucleosome structure to facilitate or prevent access of regulatory factors to the genome and are conserved throughout the eukaryotic kingdom (1-3). Early work in yeast determined that genes encoding proteins involved in mating type switching (SWI) and in sucrose fermentation (SNF) formed a multi-protein complex containing a DNA-dependent ATPase of the SNF2 family (4-6). Subsequent work in both yeast and human cells determined that such complexes altered the structure of nucleosomes in an ATP-dependent manner and promoted binding of transcription factors to the nucleosome (1, 2, 7, 8). Formal demonstration of enzymatic activity followed (5). Studies of SWI/SNF complexes in mammalian cells revealed that significant diversity exists in the assembly of the complexes (4, 9-15). There are two mutually exclusive ATPases, called Brahma (BRM) and Brahma-related gene 1 (Brg1 (9, 16)). The original purifications of mammalian SWI/SNF complexes identified two chromatographically separable fractions, originally termed “A” and “B”, that contained ATP-dependent nucleosome remodeling activity that tracked with the presence of the ATPase protein (1, 4, 8). These complexes were subsequently renamed BAF (BRG1 or BRM associated factors) and PBAF (polybromo-associated BAF), respectively, based not on the ATPase responsible for catalytic activity, but on the presence of subunits that were unique to each complex (17).

Additional work has shown that specific mSWI/SNF subunit proteins have different splice forms, can be encoded by multiple genes that are differentially expressed among tissue types or under different conditions, and consequently can be assembled into many distinct complexes that differ in subunit composition and function (18, 19). Based on the number of identified subunits and their variants, in theory there can be thousands of combinations that can constitute a mammalian SWI/SNF (mSWI/SNF) complex (18). Nevertheless, the diverse compositions of complexes could still be classified as members of the BAF or PBAF families. More recently, a third subfamily of complex has been recognized, called non-canonical BAF (ncBAF, (18)). ncBAF lacks some of the subunits that are common to BAF and PBAF and has other subunits not found in those two subfamilies (18, 20).

The catalytic activity of Brg1/Brm and homologs form other species share a common mechanism by which ATP-hydrolysis powers alteration of the path and position of the DNA around a nucleosome particle (2, 5, 21). Consequently, the diversity of mSWI/SNF complexes likely originated from the need to target the catalytic activity to different places in the genome, as well as provide specialized functions.

Research related to mSWI/SNF contributions towards skeletal muscle differentiation has been focused on the requirement and roles for the catalytic subunit. For instance, ectopic expression of catalytically inactive Brg1 or Brm subunits inhibited differentiation, prevented chromatin remodeling of several myogenic genes and consequently decreased their expression (8, 22-26). Brg1 and Brm have differential roles in regulating gene expression during myogenesis. Brg1 is required for the transcription myogenic genes at early stages of differentiation, while Brm is required for *Ccnd1* repression and cell cycle arrest that precedes the activation of late muscle genes (27). Core subunits shared by BAF, PBAF and ncBAF complexes have been also implicated in the development of heart and skeletal muscle development during mouse embryogenesis (28). Among these, the Baf60c subunit has a critical role in targeting mSWI/SNF enzymes to myogenic promoters (29). Baf53a, another core component of the three subfamilies, and Snf5, a subunit shared only by BAF and PBAF complexes, are also required for the transcriptional activation of myogenic genes (30).

mSWI/SNF complexes are also essential for myoblast proliferation. Brg1 and Snf5 subunits contribute to myoblast cell cycle progression and knockout (KO) of *Brg1* in primary myoblasts resulted in cell death (30, 31). Brg1 is essential for *Pax7* expression, a master transcription factor that supports myoblasts proliferation and survival (31). The regulation of *Pax7* expression is partially regulated by casein kinase 2-mediated phosphorylation of Brg1 (32). Moreover, short hairpin RNA (shRNA) knockdown (KD) of distinguishing subunits of the BAF, ncBAF, and PBAF complexes in C2C12 myoblasts showed the differential contributions of each subfamily to myoblast proliferation. In this regard, KD of specific subunits of the BAF or the ncBAF complexes reduced myoblast proliferation rate, while KD of PBAF-specific subunits did not affect proliferation (19). RNA-seq analyses from proliferating myoblasts KD for *Baf250A* (BAF complex) exhibited a reduction in *Pax7* expression due to a decreased binding of Baf250A and impaired chromatin remodeling at the promoter of this proliferation marker (19). The proliferation defect was reversed by reconstituting Pax7 expression using a doxycycline-inducible lentiviral vector. Thus, the work demonstrated that the BAF subfamily is required for myoblast proliferation via regulation of Pax7 expression.

In the current work, we aimed to understand the specific contributions of BAF, ncBAF and PBAF enzymes to myogenesis, using a similar strategy of shRNA-mediated KD of the the expression of the individual mSWI/SNF enzyme subunits that are specific to the three subfamilies (Baf250A, Brd9 and Baf180, respectively (19)). Our data show that the BAF complex is required for myoblast differentiation as it regulates the expression of myogenin and other myogenic genes required to initiate and continue the myogenic program. Mechanistic analyses determined that Baf250A is bound to the regulatory sequences controlling the expression of these genes. RNA-seq analyses of differentially expressed genes in *Brd9* KD differentiating myoblasts did not identify terms associated with skeletal muscle, and immunostaining of differentiating myoblasts revealed no colocalization of Brd9 and Baf250A, suggesting that the ncBAF complexes contributed to myogenesis indirectly. The PBAF subfamily was verified to be dispensable for myoblast differentiation as was previously reported (33). Our work provides functional and mechanistic details of the differential regulatory roles of the different subfamilies of mSWI/SNF enzyme complex during skeletal muscle differentiation and supports a role for the BAF complex as an essential contributor to this process.

## Materials and methods

### Antibodies

Hybridoma supernatants were obtained from the Developmental Studies Hybridoma Bank (University of Iowa) against myogenin (F5D, deposited by W. E. Wright) and anti-myosin heavy chain (MHC; MF20, deposited by D. A. Fischman). Mouse anti-Brg1 (G-7; sc-17796) and normal rabbit IgG (sc-2027) were from Santa Cruz Biotechnologies. The rabbit anti-Brd9 antibody was from Invitrogen (PA5-113488). The Mouse anti-BRD9 antibody (1H8, CBMAB-0174-YC) was from Creative Biolabs. The Rabbit anti-PBRM (Baf180, A0334), - BAF250A (A16648), -vinculin (A2752), -DPF2 (A13271), and -GAPDH (A19056) antibodies were from Abclonal Technologies. Secondary antibodies used for western blot were HRP-conjugated anti-mouse and anti-rabbit (31430 and 31460, respectively), and for immunofluorescence were the goat anti-rabbit IgG Alexa Fluor Plus 594 and the goat anti-mouse IgG Alexa Fluor Plus 488 (A32740 and A32723, respectively) that were obtained from Thermo Fisher Scientific.

### Cell culture

C2C12 and HEK293T cells were purchased from ATCC (Manassas, VA) and were maintained at sub-confluent densities in proliferation media containing Dulbecco’s modified Eagle’s medium (DMEM) supplemented with 10% fetal bovine serum (FBS) and 1% penicillin-streptomycin in a humidified incubator at 37°C with 5% CO_2_. Differentiation of C2C12 cells was initiated after cells reached 80% confluence. Cell differentiation was induced with differentiation media (DMEM supplemented with 2% horse serum, 1% Insulin-Transferrin-Selenium-A supplement (Invitrogen) and 1% penicillin-streptomycin) in a humidified incubator at 37°C with 5% CO_2_. Samples of differentiated myoblasts were processed after 48 h of induction of differentiation.

### Virus production for shRNA transduction of C2C12 cells

Mission plasmids (Sigma) encoding for two different shRNA against specific subunits of the three subfamilies of mSWI/SNF complexes BAF (Baf250A, DPF2), ncBAF (Brd9) and PBAF (Baf180) (**supplemental table 1**) were isolated by using the Pure yield plasmid midiprep system (Promega) following the manufacturer’s instructions. shRNA (15 µg) and the packing vectors pLP1 (15 µg), pLP2 (6 µg), and pSVGV (3 µg) were transfected using lipofectamine 2000 (Thermo Fisher) into HEK293T cells for lentiviral production. After 24 and 48 h the supernatants containing viral particles were collected and filtered using a 0.22 µm syringe filter (Millipore).

Proliferating C2C12 myoblasts were transduced with lentivirus in the presence of 8 µg/ml polybrene and selected with 2 µg/ml puromycin (Invitrogen).

### RT-qPCR gene expression analysis

RNA was purified from three independent biological replicates of proliferating and differentiated C2C12 myoblasts with TRIzol (Invitrogen) following the manufacturer’s instructions. cDNA synthesis was performed with 500 ng of RNA as template, random primers, and SuperScript III reverse transcriptase (Invitrogen) following the manufacturer’s protocol. Quantitative RT-PCR was performed with Fast SYBR green master mix on the ABI StepOne Plus Sequence Detection System (Applied Biosystems, Foster City, CA) using the primers listed in **supplemental table 2**. The delta threshold cycle value (ΔC_T_) was calculated for each gene and represents the difference between the CT value of the gene of interest and that of the *Eef1A1* reference gene.

### RNA-sequencing analysis

Duplicate samples for RNA sequencing were purified from differentiating C2C12 myoblasts with TRIzol (Invitrogen) following the manufacturer’s instructions. Sample quality and concentration was determined at the Molecular Biology Core Lab at the University of Massachusetts Chan Medical School, Fragment Analyzer services. RNA library preparation and sequencing was performed by the BGI Americas Corporation (Cambridge, MA, US). Briefly, libraries were sequenced using the BGISEQ-500 platform and reads were filtered to remove adaptor-polluted, low quality and high content of unknown base (N) reads. About 99% raw reads were filtered out as clean reads, which were then mapped to mouse reference genome mm10 using HISAT. Transcripts were reconstructed using StringTie (34) and novel transcripts were identified using Cufflinks (35). All transcripts were then combined together and mapped to mm10 reference transcriptome using Bowtie2 (36). Gene expression levels were calculated using RSEM (37). DEseq2 (38) and PoissonDis (39) algorithms were used to detect differentially expressed genes (DEG). GO analysis using DAVID (https://david.ncifcrf.gov/tools.jsp) was performed on DEGs to cluster genes into function-based and pathway-based categories.

### Western blot analyses

C2C12 myoblasts were washed with PBS and solubilized with RIPA buffer (10 mM piperazine-N,N-bis(2-ethanesulfonic acid), pH 7.4, 150 mM NaCl, 2 mM ethylenediamine-tetraacetic acid (EDTA), 1% Triton X-100, 0.5% sodium deoxycholate, and 10% glycerol) containing protease inhibitor cocktail (Thermo Fisher Scientific). Protein content was determined by Bradford (40). Samples (20 µg) were prepared for SDS-PAGE by boiling in Laemmli buffer. The resolved proteins were electro-transferred to PVDF membranes (Millipore). The proteins of interest were detected with the specific antibodies as indicated in the figure legends and above, followed by species-specific peroxidase conjugated secondary antibodies and chemiluminescent detection (Tanon, Abclonal Technologies).

### Immunofluorescence

C2C12 cells grown on coverslips were fixed with 3% paraformaldehyde, permeabilized with 0.5% Triton X-100, and blocked with PBS containing 1% bovine serum albumin (BSA). Primary and secondary antibodies were diluted in 1% blocking solution. The primary antibodies against Baf250A and Brd9 were used at a dilution of 1:100. Alexa Fluor-conjugated secondary antibodies (1:500 dilution) were incubated for 2 h at room temperature. Nuclei were detected with DAPI (4’,6-diamidino-2-phenylindole). Cover slides were mounted with ProLong Gold antifade reagent (Life Technologies) and imaged with inverted fluorescence microscopy (Leica DMI6000). Leica LAS AF Lite software was used for image processing.

### Immunocytochemistry analyses

C2C12 myoblasts (control and KDs) were fixed overnight in 10% formalin-PBS at 4 ºC. Cells were washed with PBS and permeabilized for 10 min with PBS containing 0.2% Triton X-100. Immunocytochemistry was performed using the hybridoma supernatants from the Developmental Studies Hybridoma Bank (University of Iowa) against myogenin and MHC. Samples were developed with the universal ABC kit (Vector Labs) following the manufacturer’s protocol.

### Calculation of fusion index for the myotubes

The fusion index was calculated as the ratio of the nuclei number in C2C12 myocytes with two or more nuclei *vs*. the total number of nuclei as previously described (41). Edges and regions that did not show good cell adhesion were discarded from the analysis. Three independent biological replicates were grown in 24-well plates and cells were induced to differentiate as described above. Quantitative analysis was performed using ImageJ software v.1.8 ((42, 43) National Institutes of Health, Bethesda, MD, USA).

### Chromatin immunoprecipitation assays

Chromatin immunoprecipitation assays were performed as previously described (20, 91). Briefly, differentiating (48 h) C2C12 myoblasts were cross-linked with 1% formaldehyde (Ted Pella Inc.) for 10 min at room temperature. Formaldehyde quenching was done with 125 mM glycine for 5 min. Crosslinked myoblasts were washed twice with ice-cold phosphate-buffered saline (PBS) supplemented with protease inhibitor cocktail and lysed with 1 ml of ice-cold buffer A (10 mM Tris HCl [pH 7.5], 10 mM NaCl, 0.5% NP-40, 0.5 µM dithiothreitol [DTT], and protease inhibitors) by incubation on ice for 10 min. The nuclei were pelleted by centrifugation at 3000 x *g*, washed with 1 ml of buffer B (20 mM Tris HCl [pH 8.1], 15 mM NaCl, 60 mM KCl, 1 mM CaCl_2_, 0.5 µM DTT). DNA was sheared by incubating the nuclei in 100 µl of buffer B supplemented with 1,000 units of micrococcal nuclease (M0247S; NEB) for 30 min at 37°C; the reaction was stopped by adding 5 µl of 0.5 M EDTA. Nuclei were pelleted and resuspended in 400 µl of ChIP buffer (100 mM Tris HCl [pH 8.1], 20 mM EDTA, 200 mM NaCl, 0.2% sodium deoxycholate, 2% Triton X-100, and protease inhibitors), and sonicated for 10 min (medium intensity, 30 s on/30 s off) in a Bioruptor UCD-200 system (Diagenode), and centrifuged at 21,000 x *g* for 5 min. The length of the fragmented chromatin was between 200 and 500 bp as analyzed on agarose gels. ChIP was performed by incubating specific antibodies against Brg1, Brd9, Baf170, Dpf2, or Baf250A with each sample for 2 h at 4 ºC. Anti-IgG ChIPs were included as negative controls. Immunocomplexes were recovered with 20 µl of magnetic Dynabeads (Thermo Fisher Scientific) after an overnight incubation at 4°C. Three sequential washes with low salt ChIP buffer followed with one final high salt washing step were performed to eliminate unspecific binding. Complexes were eluted in 100 µl of elution buffer (0.1 M NaHCO_3_, 1% SDS) for 30 min at 65°C, incubated with 1 µl of RNAse (0.5 mg/ml) for 30 min at 37°C, and reverse cross-linked by addition of 6 µl of 5M NaCl and 1 µl of proteinase K (1 mg/ml) overnight at 65°C. DNA was purified using a ChIP DNA Clean & Concentrator kit (Zymo Research). Bound DNA fragments were analyzed by quantitative PCR using SYBR green master mix. Quantification was performed using the fold enrichment threshold cycle method 2^Δ(CT sample – CT IgG)^, and data are shown relative to the results determined for the IgG control. The primer sequences are listed in **supplemental table 1**.

### Statistical analysis

Statistical analysis was performed using Kaleidagraph (Version 4.1) or Graph Pad Prism 7.0b. Multiple data point comparisons and statistical significance were determined using one-way analysis of variance (ANOVA). Experiments where p<0.05 were considered to be statistically significant.

## Results

### The BAF subfamily of mSWI/SNF complexes is required for skeletal muscle differentiation *in vitro*

Evaluation of proliferating primary and immortalized C2C12 myoblasts identified Brg1 as the relevant mSWI/SNF ATPase and the BAF subfamily of mSWI/SNF complexes as the major contributors to maintenance of the proliferative state by regulating the expression of *Pax7* (19, 31, 44). We continued our work to determine the roles of the three families of mSWI/SNF complex in the differentiation of myoblasts *in vitro*. We used a lentiviral system to KD specific subunit proteins that are unique for each of the different subfamilies of mSWI/SNF complexes and induced the myoblasts to differentiate. Western blot analyses showed that differentiating C2C12 myoblasts transduced with lentiviral particles containing shRNAs against *BAF250A, Brd9*, and *Baf180*, which encode subunits that are unique to the BAF, ncBAF, and PBAF complexes, respectively, showed reduced expression of these proteins (**Fig. 1**). We then assessed the functional effects of these KDs on myoblast differentiation using changes in fusion index as a quantitative measure of the progression of myogenesis. **Figure 2** and **Supp. Fig. 1** show representative micrographs of control and KD C2C12 myoblasts undergoing differentiation for 24, 48, 72 and 96 h. Myogenin and MHC expression in wild type (untreated) and scrambled sequence shRNA (scr) controls were detected at 24 h after inducing differentiation; myoblast fusion was detected as early as 48 h and peaked at 96 h (**Fig. 2A, Supp. Fig. 1A**). Myoblasts partially depleted of Baf250A (BAF complex; **Fig. 2B, Supp. Fig. 1B**) failed to differentiate as shown by a decrease in the expression of myogenin and MHC and a reduction in the fusion index when compared to control cells at similar time points (**Fig. 2A, Supp. Fig. 1A**). The KD of the Brd9 subunit unique to the ncBAF complex (**Fig. 2C, Supp. Fig. 1C**) resulted in a delay in the differentiation progression, as shown by a decrease in expression of differentiation markers and longer times required for these myoblasts to fuse. However, the Brd9 KD myoblasts were able to differentiate within the 96 h analyzed (**Fig. 2C, Supp. Fig. 1C**). Partial depletion of Baf180 subunit that is unique to the PBAF complex had no effect on myogenesis, confirming previous results of our group and others that showed that it is dispensable for skeletal muscle differentiation (**Fig. 2D, Supp. Fig. 1D;** (33, 45)). The data supports a fundamental role for the BAF complex in myoblast differentiation.

**Figure 1.**
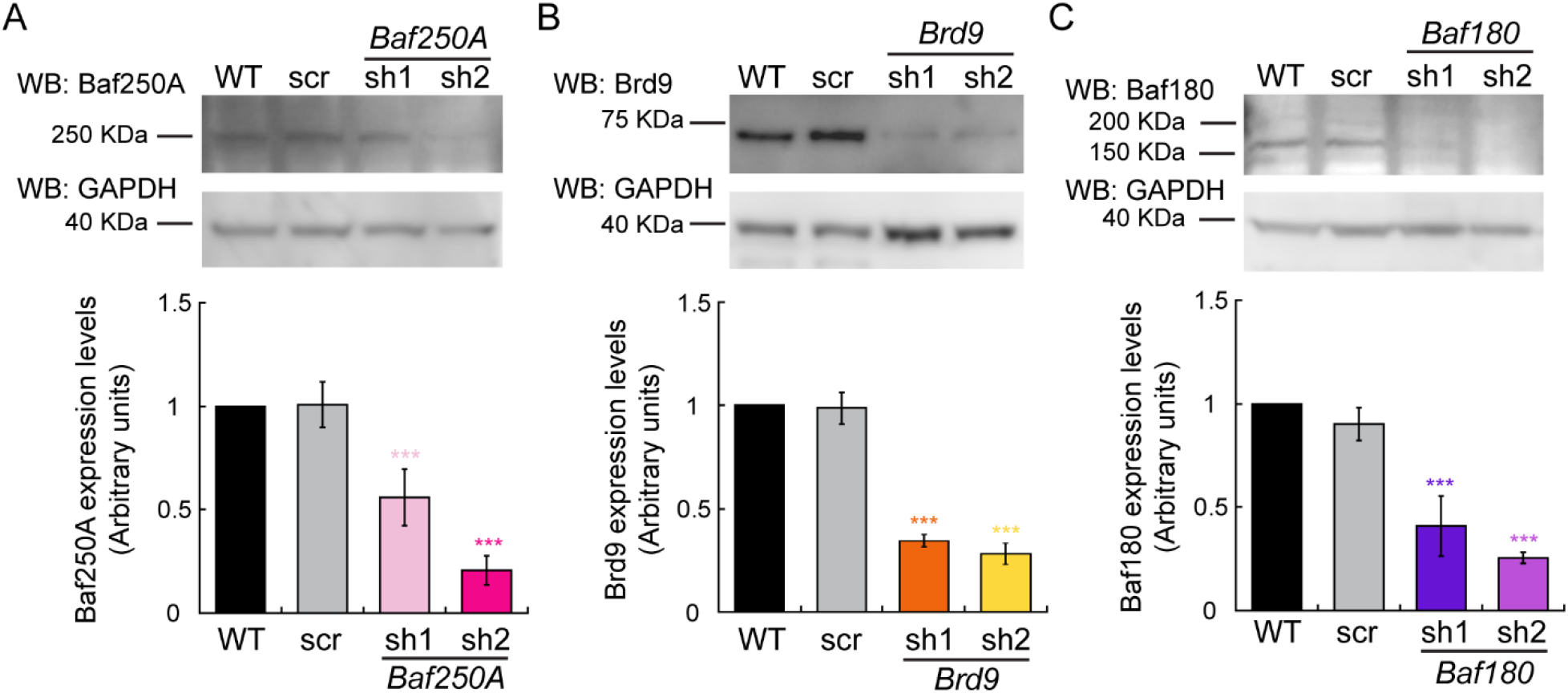
Baf250A, Brd9 and Baf180 expression in wild type (WT), shRNA scrambled (scr) control, and the indicated shRNA-mediated knockdowns in differentiating C2C12 myoblasts. Representative western (top) and quantification (bottom) of Baf250A (**A**), Brd9 (**B**), and Baf180 (**C**) levels in differentiating cells after 48 h of differentiation. Data represent the mean ± SE of three independent biological replicates. GAPDH was used as the loading control. Quantification of each sample was compared to the corresponding wild type (WT) sample, which was set to 1.0. ***P < 0.001.

**Fig. 2.**
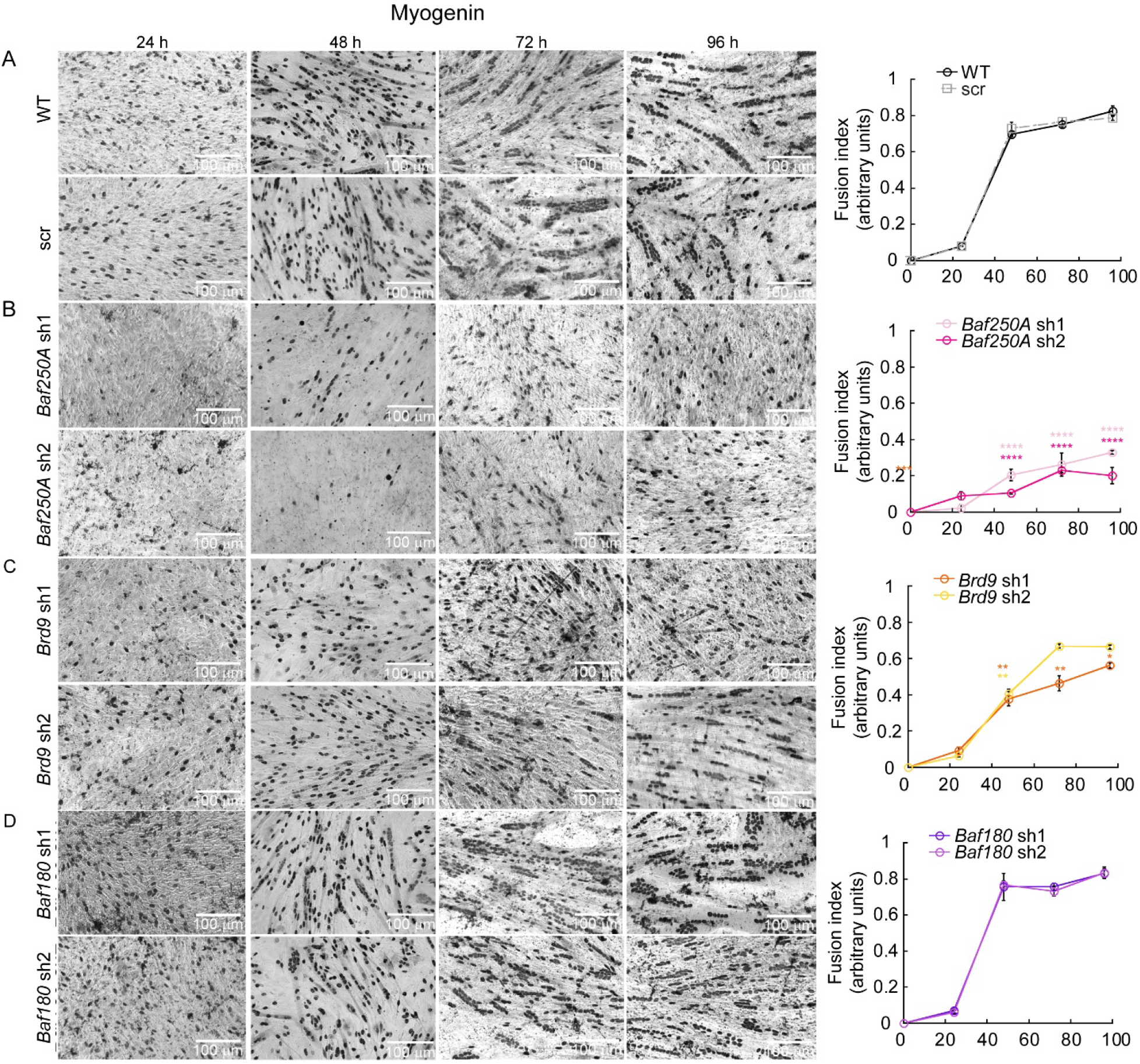
Baf250A knockdown inhibited the differentiation of C2C12 cells. Representative light micrographs and fusion index of wild type C2C12 myoblasts or myoblasts transduced with either scr, *Baf250A, Brd9* or *Baf180* shRNAs undergoing differentiation for 24, 48, 72 and 96 h Cells were immunostained for Myogenin. Bars = 100 µm. *P < 0.05, **P < 0.01, ****P < 0.0001.

### KD of *BAF250A* or *BRD9* elicited differential effects in gene transcription in differentiating C2C12 myoblasts

We have previously shown that KD of subunits unique to the three different subfamilies of the mSWI/SNF complexes had minor effects on gene expression in proliferating myoblasts (19). In that report, one of the most striking results was the effect of *BAF250A* KD in the expression of *Pax7*, the master regulator of myoblast growth, which partially explained the growth deficiency observed in those cells (19). KD of BAF250A also impaired myoblast differentiation, suggesting a primary role of the BAF complex in this process. To better understand the phenotypes observed in myogenesis at a transcriptional level, we performed RNA-seq analyses in differentiating myoblasts transduced with scramble (Scr) shRNA, and myoblast expressing shRNAs against *Baf250A, Brd9* or *Baf180* (**Fig. 3**). In all cases, the sequenced libraries had approximately 45 M total reads. Pearson coefficients were >0.96 for the replicate RNA-seq datasets for each KD and are shown in **Supplemental Table 3**. The data were mapped to the mouse genome (mm10), and changes in gene expression were determined. **Supplemental Table 4** shows the differentially expressed genes that showed significant changes in both replicates for each shRNA (log2FoldChange < 1, padj < 0.05). The *Baf250A* KD resulted changes in the expression of 4729 genes when compared to control cells; of these, 1844 genes were upregulated and 2885 were downregulated. (**Fig. 3A, Supplemental Table 4**). *Brd9* KD affected 2483 genes of which 879 were upregulated and 1604 downregulated (**Fig. 3B, Supplemental Table 4**). In addition, we determined there were 2052 genes that were altered in both BAF250A and BRD9 KD cells when compared to control, which represents ∼83% of the genes differentially regulated by BRD9. In contrast, only ∼43% of the differentially regulated genes in BAF250A KD cells are shared with those genes differentially regulated by BRD9 KD (**Fig. 3C**). These results suggest that there are relatively few genes uniquely regulated by Brd9 in differentiating myoblasts.

**Fig. 3.**
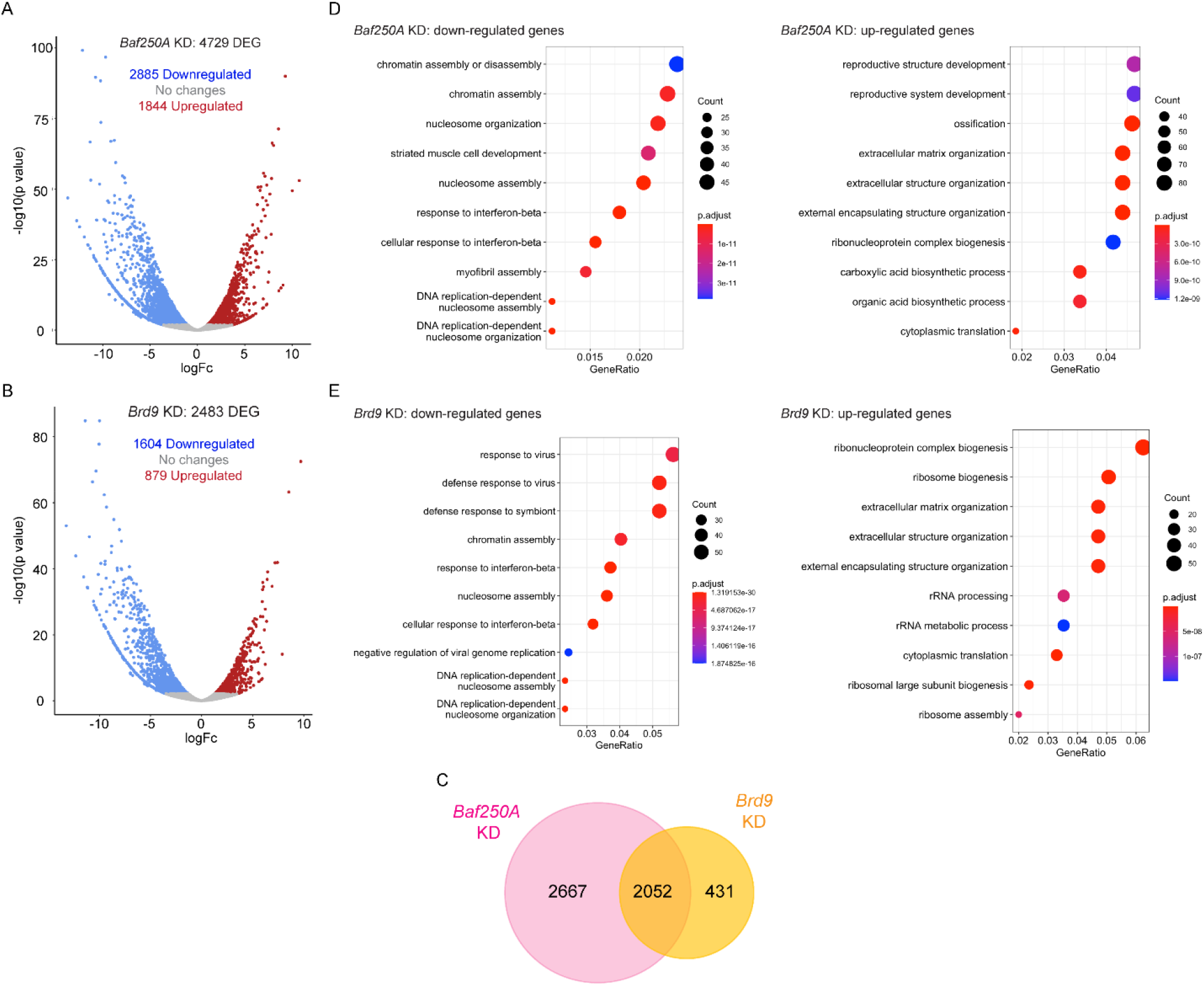
Changes in gene expression dependent on Baf250A or Brd9 knockdown. Volcano plots displaying differentially expressed genes between *scr* control and *Baf250A* **(A)**, or *Brd9* **(B)** knockdown C2C12 cells. The y-axis corresponds to the mean log10 expression levels (P values). The red and blue dots represent the up- and down-regulated transcripts in knockdown cells (false-discovery rate [FDR] of <0.05), respectively. The gray dots represent the expression levels of transcripts that did not reach statistical significance (FDR of >0.05). (**C)** Venn diagram showing the overlapping differentially expressed genes between the differentiating myoblasts KD for the *Baf205A* and *Brd9* subunits. GO term analysis of differentially expressed genes in differentiating C2C12 cells KD for *Baf250A* **(D)** or *Brd9* **(E)**. Cut-off was set at 2.0 of the -log(adjusted P value). See Supp. Table 4 for the complete list of genes.

Gene ontology (GO) analysis of these differentially expressed genes identified functional categories. GO analyses of *Baf250A* KD differentiating myoblasts determined a deficiency in the expression of genes related to chromatin regulation and nucleosome assembly, which would be expected if a chromatin remodeling enzyme was compromised, response to interferon-β, and muscle development and structure (**Fig. 3D**). Upregulated genes were related to reproductive system development, ossification, and extracellular matrix organization (**Fig. 3D**). Increases in bone-specific gene expression upon inhibition of muscle-specific gene expression is expected (46). Genes involved in ECM and the extracellular environment are known to be regulated by Brg1 (47, 48). *Brd9* KD myoblasts showed decreased expression of genes involved in responses to virus, response to interferon-β signaling and chromatin regulation, similar to the results from the *Baf250A* KD (**Fig. 3E**). Notably, terms related to muscle formation or function were absent, suggesting that only Baf250A, and by extension, the BAF complex, regulates muscle-specific genes. The top terms describing genes with enhanced expression upon Brd9 KD were related to ribonucleoprotein and ribosomal synthesis and extracellular matrix organization (**Fig. 4E**), demonstrating some overlap with the terms upregulated in response to Baf250A KD.

**Fig. 4.**
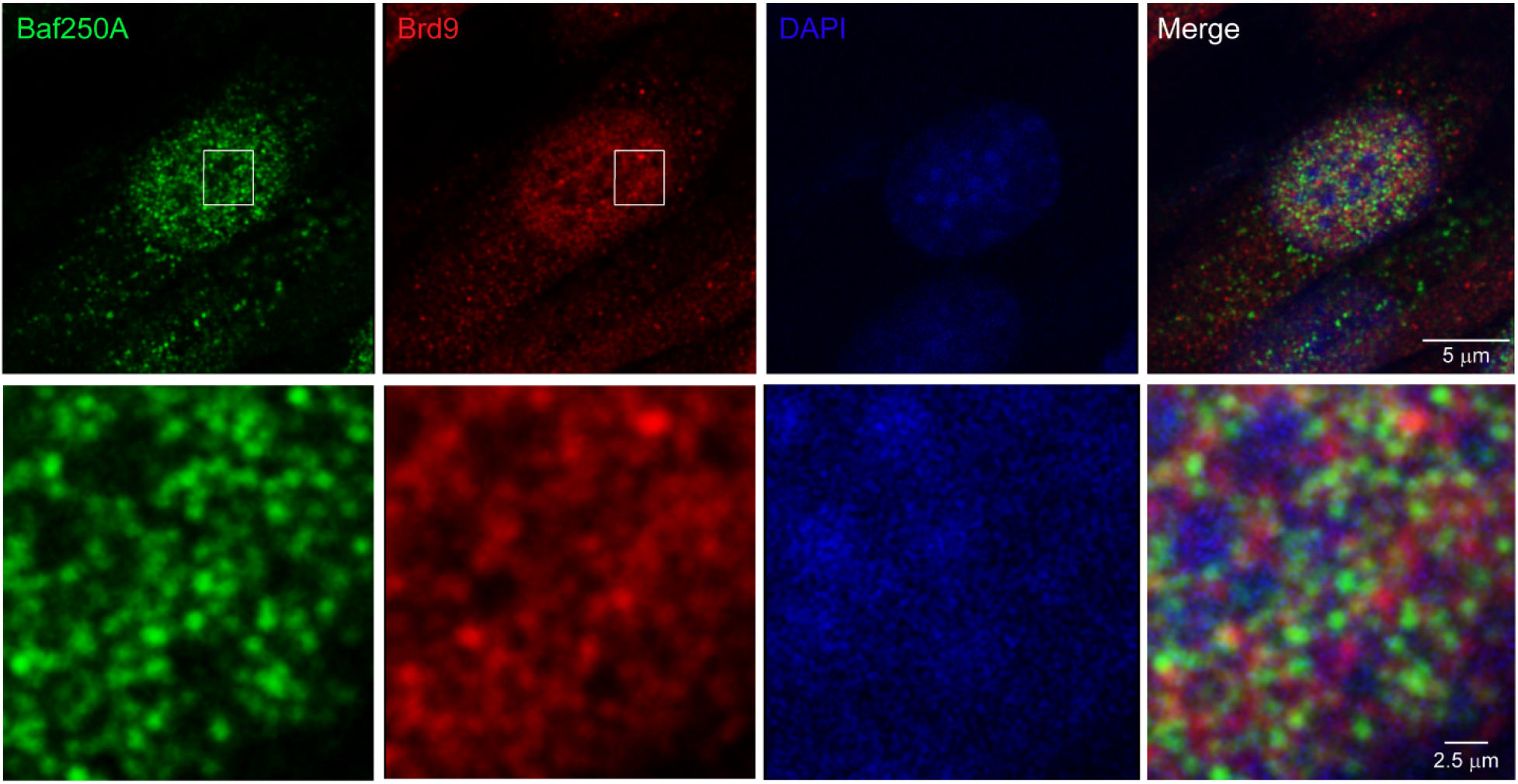
Baf250A and Brd9 do not colocalize in differentiating C2C12 myoblasts. Representative confocal images showing the nuclear localization of Baf250A (green) and Brd9 (red) in differentiating (48 h) wild type C2C12 myoblasts. Zoom in area is delimited with a square in the top images and depicted in bottom panel. The nuclei were stained with DAPI (blue). Bars = 5 and 2.5 μm, respectively.

### BAF250A and BRD9 do not colocalize in differentiating myoblasts

The RNA-seq data suggested that muscle-specific gene expression was regulated by Baf250A but not by Brd9. We examined nuclear localization for Baf250A and Brd9 and determined that there was essentially no overlap in the localization of these proteins (**Fig. 4**). This result strongly suggests that Baf250A and Brd9 are spatially separated in the cell nucleus and are therefore unlikely to be acting together at the same DNA sequence. This observation reinforces the idea that Baf250A, and not Brd9, mediates control of myogenic genes. It also suggests that genes that are similarly affected by Baf250A and Brd9 can be regulated by either and are unlikely to be regulated by both at the same time.

### BAF complex-mediated regulation of myogenic genes

Dpf2, also called Baf45D, is another mSWI/SNF subunit that, like Baf250A, is specific to the BAF complex. shRNA-mediated KD of *Dpf2* in C2C12 cells was achieved (**Supp. Fig. 2**); these cells were compromised for myogenic differentiation as judged by MHC staining and fusion index determinations (**Supp. Fig. 3**), mirroring what was observed upon *Baf250A* KD.

We then validated expression of a few myogenic genes by quantitative reverse transcriptase PCR (qRT-PCR), in cells KD for Baf250A or Dpf2. We quantified steady-state mRNA levels of the differentiation markers myogenin (*Myog*), muscle specific creatine kinase (*Ckm*), caveolin 3 (*Cav3*) and myosin heavy chain IIb (*MyHCII*). We detected a significant decrease in the amount of each myogenic transcript in cells KD for Baf250A or Dpf2 (**Fig. 5**).

**Fig. 5.**
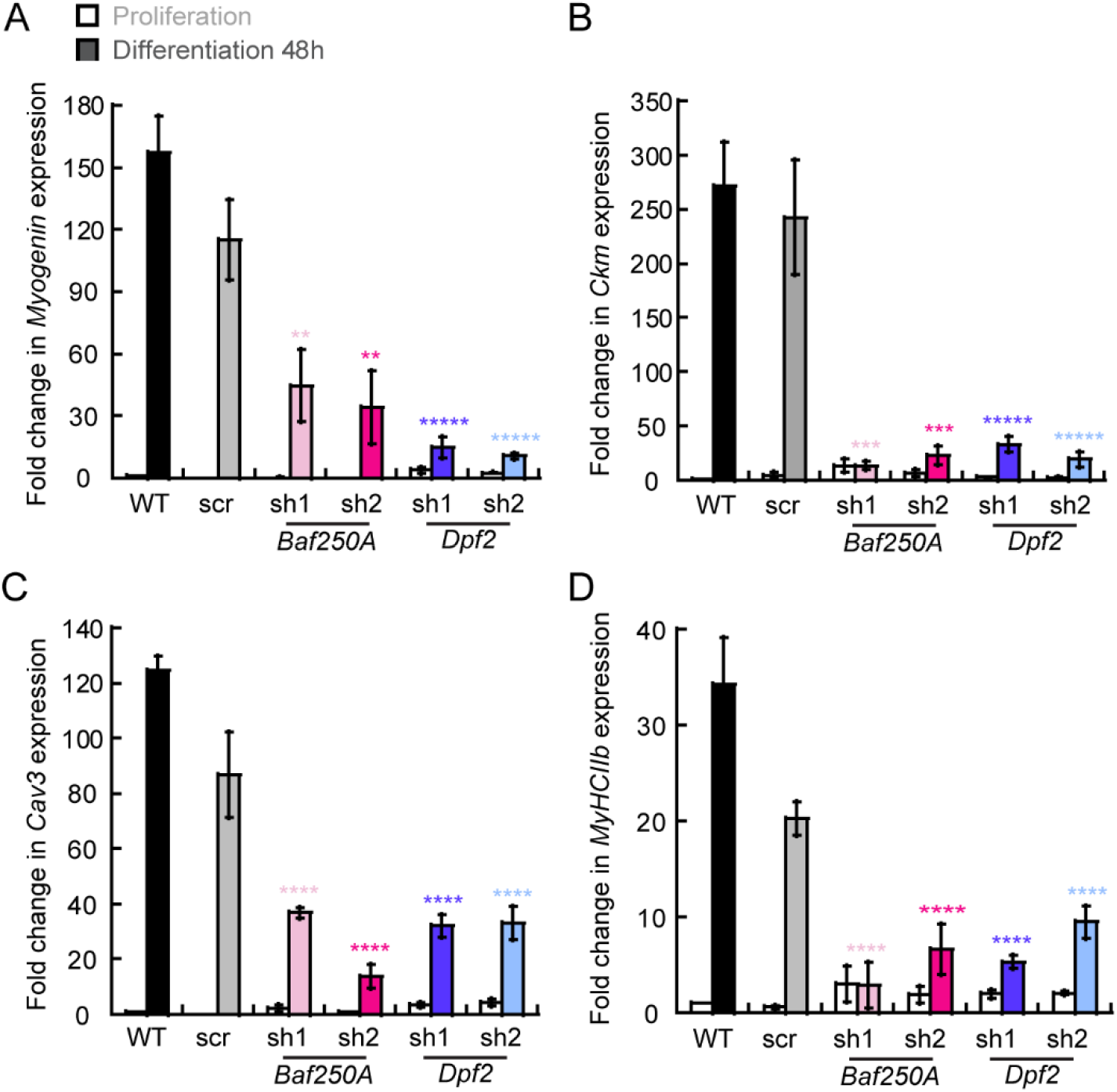
The expression of myogenic genes is impaired in differentiating myoblasts KD for the BAF complex subunits *Baf250A* and *Dpf2*. Steady state mRNA levels of (**A**) *Myogenin*, (**B**) muscle-specific creatine kinase (*Ckm*), (**C**) Caveolin 3 (*Cav3*), and (**D**) myosin heavy chain II (*MhcII*) determined by qRT-PCR from proliferating (white bars) and differentiating (filled, colored bars) C2C12 myoblasts. Cells were transduced with either *scr, Baf250A* or *Dpf2* shRNAs. For each gene the data were normalized against expression in proliferating control cells, which was set at 1.0, and represent the mean ± SE for three independent experiments. * P < 0.05, ** P < 0.01, *** P < 0.001, **** P < 0.0001.

It is well-established that the mSWI/SNF ATPases can be localized by ChIP methods to regulatory sequences controlling the expression of myogenic genes and that knockdown or inhibitors of the bromodomains located in the ATPase protein structure can inhibit binding to target sequences (33). Since the BAF complex is required for myoblast differentiation and activation of myogenic gene expression, we would predict that KD of BAF complex-specific subunits would deleteriously impact binding of the BAF enzyme to myogenic gene regulatory sequences. The data showed that there was a significant decrease in *Baf250A* binding to myogenic gene regulatory sequences in cells partially depleted of this subunit (**Fig. 6A**). KD of the *Dpf2* subunit similarly decreased binding to the myogenic gene regulatory sequences tested (**Fig. 6B**).

**Fig. 6.**
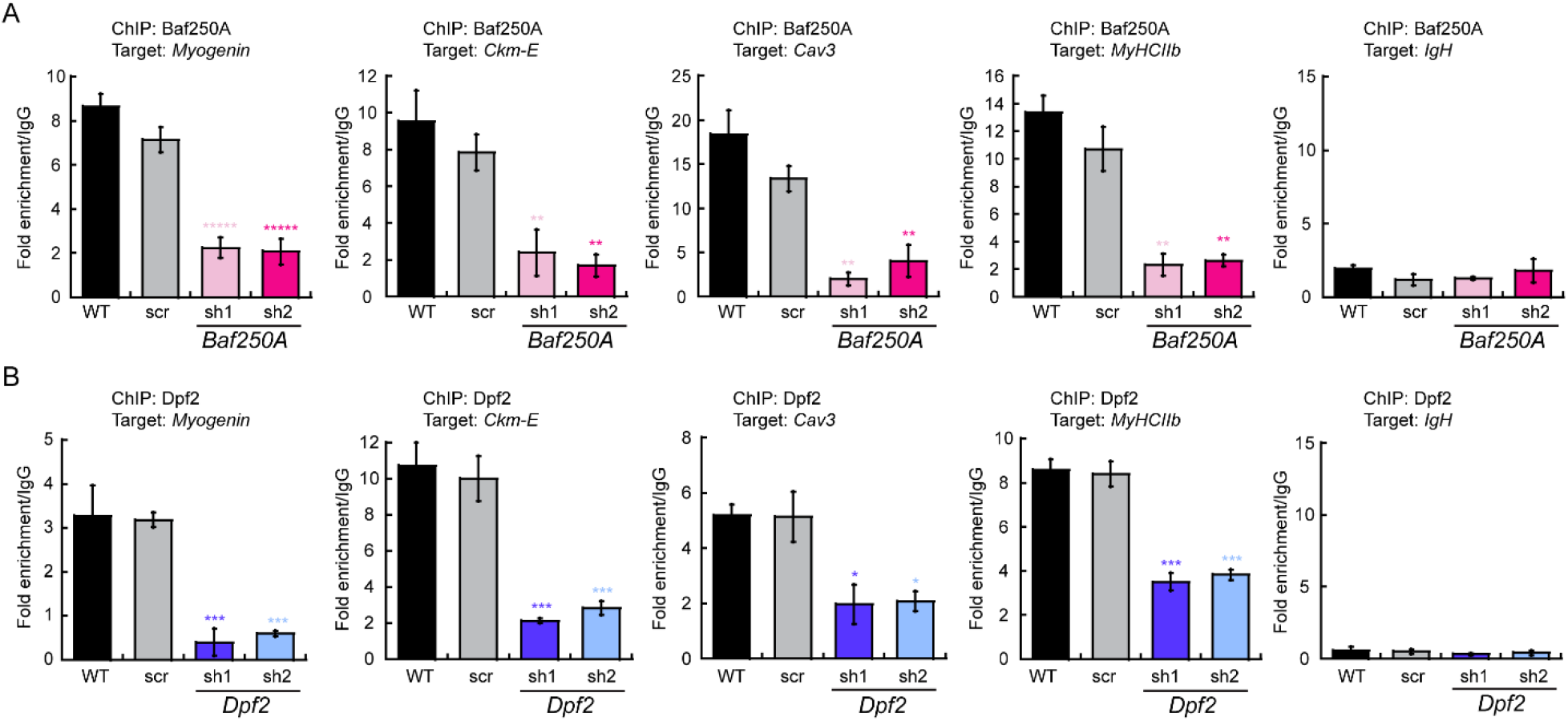
Effect of KD of BAF complex components in the binding of specific subunits to the *Myogenin* promoter. ChIP-qPCR showing binding of Baf250A (**A**) and DPF2 (**B**) to the *Myogenin* promoter, the muscle creatine kinase enhancer (*Ckm-E*), Caveolin 3 (*Cav3*), myosin heavy chain II (*MhcII*), and immunoglobulin H (*IgH*; negative control) in 48 h differentiating C2C12 myoblasts. C2C12 myoblasts were transduced with either *scr, Baf250A* or *Dpf2* shRNAs. Data are the mean ± SE for three independent experiments. * P < 0.05, ** P < 0.01, *** P < 0.001, ***** P < 0.00001.

Prior work of how mSWI/SNF complexes belonging to different subfamilies assemble in solution has determined that the BAF-specific subunits, Baf250A and Dpf2, join after the “core” subunits are assembled into a sub-complex. The ATPase subunits join subsequently and are among the last of the subunits to join the complex (18). We asked whether KD of Baf250A or of Dpf2 would impact the binding of the Brg1 ATPase that is critical for catalytic function. As expected, Brg1 binding was compromised if Baf250A or Dpf2 levels were reduced by KD (**Fig. 7A**). In contrast, the Baf170 subunit is required to initiate the formation of the core complex that can exist in the absence of Baf250A or Dpf2 (18). We asked whether or not Baf170 was bound to myogenic regulatory sequences in the presence of Baf250A or Dpf2 KD. We repeated the ChIP analysis for Baf170 and determined that its binding was also compromised at the target sequences (**Fig. 7B**). KD of either BAF-specific subunit therefore compromises the ability of the BAF complex to interact with target sequences and does not support the idea that a partial BAF complex can stably bind.

**Fig. 7.**
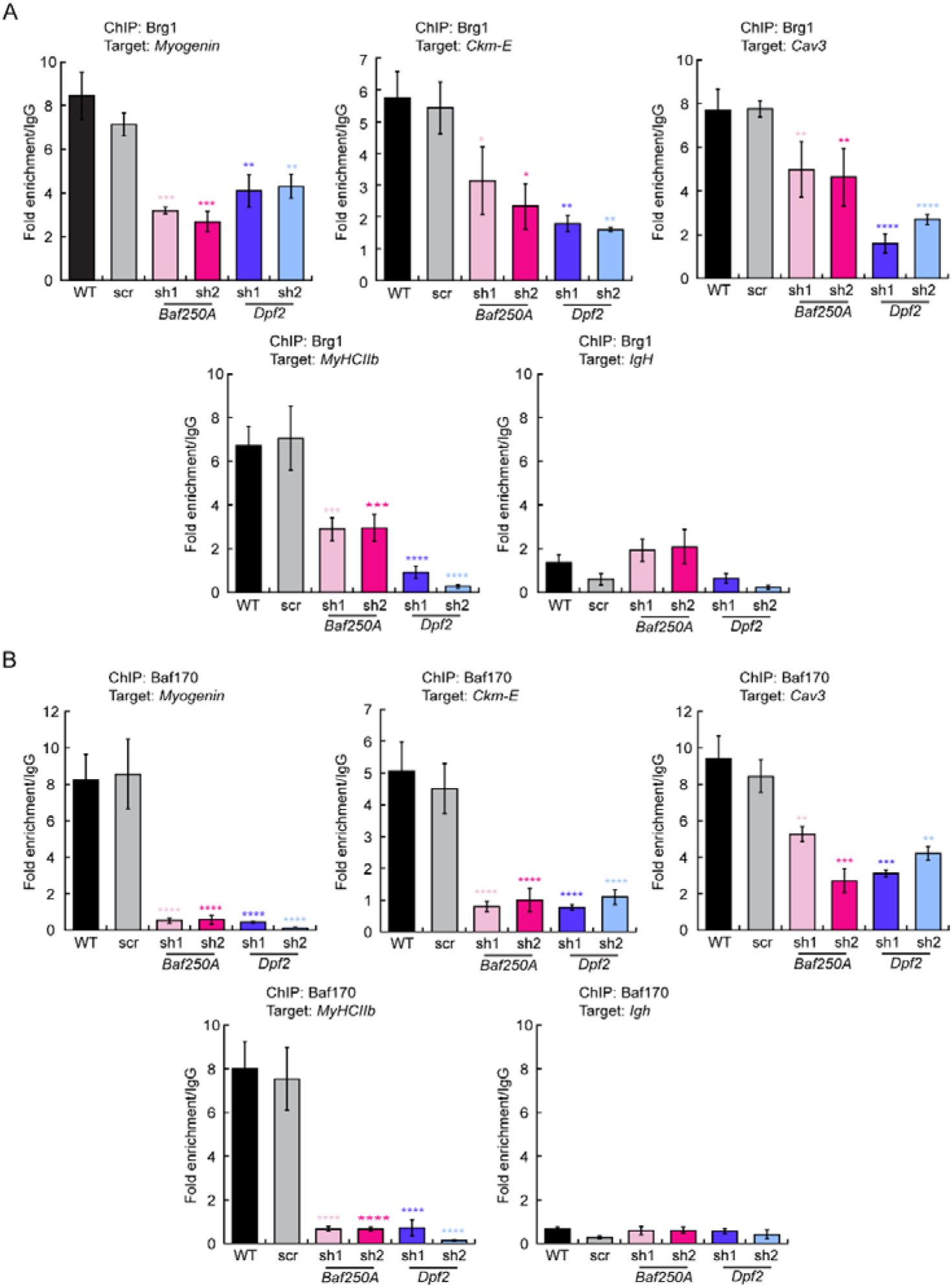
Binding of Brg1 and Baf170 to *myogenic gene regulatory sequences* is compromised upon KD of *Baf250A* or *Dpf2*. ChIP-qPCR showing binding to the indicated sequences of **(A)** Brg1 and **(B)** Baf170 in differentiating C2C12 myoblasts that were transduced with either *scr* or *Baf250A* or *Dpf2*. Binding in untransduced, differentiating C2C12 cells was shown for comparison. Binding to the *IgH* gene is shown as a negative control. Data are the mean ± SE for three independent experiments. * P < 0.05, ** P < 0.01, *** P < 0.001, **** P < 0.0001.

## Discussion

Cell and tissue lineage determination and differentiation are complex processes that rely on the tight regulation of gene expression by transcription factors, chromatin remodeling enzymes, and other co-regulatory proteins. Transcriptional regulators can be expressed in a tissue-specific manner to modulate the expression of tissue-specific genes to allow cells to differentiate into tissues and organs and to enable the growth and development of eukaryotes (49, 50). Among these proteins, chromatin remodelers of the mSWI/SNF family drive developmental events that enable cells to grow and differentiate (51-55). These enzymes utilize the energy released from ATP hydrolysis to modify the structure of nucleosome and the chromatin environment at specific loci, depending on the cellular requirements (56-58). The mSWI/SNF complex have been divided in three different families based on their subunit composition though they are sometimes separated by the presence of either of the catalytic subunits, BRG1 or BRM (4, 7, 18, 59).

In this work, we expanded our studies on the specific biological roles of the three subfamilies of the mSWI/SNF complex by examining myoblast differentiation. Myogenesis is characterized by the expression of myogenic regulatory factors (MyoD, MRF4, MYF5 and Myogenin) that bind to consensus E-boxes at the promoters and enhancers of myogenic genes (56, 60-62). These cooperate with the family of myocyte enhancer factor 2 (MEF2) proteins and other transcription factors to promote the expression of a downstream cascade of myogenic genes (63, 64). Recruitment of the mSWI/SNF enzymes by these transcription factors is required for activation of myogenic genes expressed both early and late in the differentiation process (8, 23-25, 27, 29).

Here, we demonstrated that the BAF subfamily of mSWI/SNF enzymes, physically defined by the presence of the Baf250A and Dpf2 subunits, specifically mediates activation of the myogenic gene expression program. Knockdown of either protein inhibited myoblast differentiation, expression of representative myogenic marker genes, and binding of not just those subunits, but other mSWI/SNF subunits at myogenic gene regulatory sequences. Go term analysis of genes that were down-regulated in differentiating myoblasts subjected to Baf250A KD identified categories related to muscle development and structure, providing corroborating evidence that the BAF subfamily of mSWI/SNF enzymes contributes to myoblast differentiation. Our prior studies suggested that the BAF subfamily was predominantly responsible for myoblast proliferation and for expression of the *Pax7* regulator of myogenic proliferation and viability.

This suggests an essential role for the BAF subfamily from at least the point of myoblast specification through myoblast formation.

In contrast, KD of the Brd9 subunit, which is specific to the ncBAF complex, had a much more modest effect on myoblast differentiation, and analysis of RNA-seq data did not identify terms related to muscle differentiation and function, suggesting that the impact of ncBAF complex disruption on myoblast differentiation is indirect. Immunofluorescence studies demonstrating a lack of co-localization of Baf250A and Brd9 in differentiating myoblasts further support this conclusion. This observation indicates that the BAF and the ncBAF subfamilies of enzymes are not in the same location, suggesting that they do not regulate the same target sequence at the same time. It will be interesting to determine if the observed spatial separation of these mSWI/SNF subfamilies is a general phenomenon or occurs under specific biological circumstances. We speculate that the effects of Brd9 KD on myoblast differentiation are due to dysregulation of ribosomal biogenesis and rRNA processing, as seven of the top ten GO terms identified from the pool of genes upregulated in Brd9 KD cells related to rRNA, ribosomes, and translation. mSWI/SNF proteins recently have been linked to altered translational efficiency in cancer cells via direct mechanisms and via inhibition of mSWI/SNF protein function by chemical inhibitors (33); it is possible that altering the gene expression and the processing of ribosome components also promotes altered translational efficiency.

The Baf180 protein, which is unique to the PBAF subfamily of mSWI/SNF enzymes, was not required for myoblast differentiation. The work here complements prior cell culture and in vivo studies indicating Baf180 is dispensable for myogenesis (33, 45). Prior work also demonstrated that Baf180, and by extension, PBAF complex, was dispensable for myoblast proliferation and for activation of the *Pax7* regulator of myoblast proliferation (19).

There is a paucity of information on the functional distinctions between the different subfamilies of mSWI/SNF enzymes in development. We have presented evidence that the BAF subfamily is primarily responsible for maintaining myoblasts in the proliferative state via activation of the *Pax7* gene (REF) and for initiating lineage-specific gene expression during differentiation. Baf250A is also required for cardiac precursor cell differentiation and for proper heart formation (65-68). However, PBAF-specific subunits, including Baf180 and Baf200, are also required for heart formation (69-71) suggesting that there are either separable requirements for the two subfamilies or there is an uncharacterized mechanism of cooperation. Baf250A has also been implicated in neural stem cell proliferation and differentiation during cortical development (72), early embryo development, and ES cell differentiation (73-75). However, the requirement for other mSWI/SNF subfamilies was not determined for these processes. ncBAF was reported to regulate pluripotency in mouse ES cells (76), but little else is known outside the context of cancer. At present, general conclusions about the roles of the different subfamilies are not possible.

Studies addressing the mechanism of action of the mSWI/SNF chromatin remodelers suggest there may be stepwise assembly of the enzyme at target sequences. For instance, in proliferating myoblasts the p38 kinase phosphorylates the BAF60c subunit of the chromatin remodeler and enables its association with MYOD and recruitment to myogenic gene promoters (25, 29). The phosphorylated BAF60c-MYOD complex act as a scaffold that brings additional subunits of the mSWI/SNF complex to the myogenic loci to make chromatin accessible (29). Later studies by Mashtalir *et al*., demonstrated that the components of the three different families are recruited sequentially and that the catalytic subunit is among the last components to associate in order to ultimately modulate gene expression programs (18). Whether this programmed and organized association of the chromatin remodeler components is maintained on chromatin and/or in different tissues remains to be elucidated. However, results from this study and our prior studies suggest that knockdown of Baf250A or of Brg1, or inhibition of Brg1 with the bromodomain inhibitor PFI-3 or the calcineurin inhibitor FK506, inhibits the binding of other mSWI/SNF subunits (19, 33, 77, 78), suggesting that stable subcomplexes of different configurations of mSWI/SNF enzymes with chromatin are not common. Additional work will be needed to distinguish between the possibility of step-wise assembly of mSWI/SNF enzymes on chromatin and binding of a pre-formed complex.

## Supporting information

Padilla-Benavides Supplementary information

Padilla-Benavides Supplemental table 4

## DATA AVAILABILITY STATEMENT

The datasets presented in this study can be found in online repositories upon acceptance of the article. RNA-seq datasets are available at GEO. The accession number is: GSE196281.

## AUTHOR CONTRIBUTIONS

A.N.I. and T.P.-B. designed research, contributed reagents and funding; T.P.-B., M.O.F., H.W., and K.X. performed experiments and compiled data; T.P.-B., M.O.F., A.N.I., S.A.S and T.S. analyzed data; T.P.-B. and T.S. prepared figures and tables T.P.-B. wrote the paper the first draft of the manuscript. All authors edited and revised the manuscript; all authors approved the final version of the manuscript.

## FUNDING

This work was funded by NIH grant R35GM136393 to A.N.I. and R01AR077578 to T.P.-B.

## Notes

### Competing Interest Statement

The authors have declared no competing interest.

