## Supplementary material for "Differential requirements for different subfamilies of the mammalian SWI/SNF chromatin remodeling enzymes in myoblast differentiation": Padilla-Benavides Supplementary information

**Supplementary Figure 1**

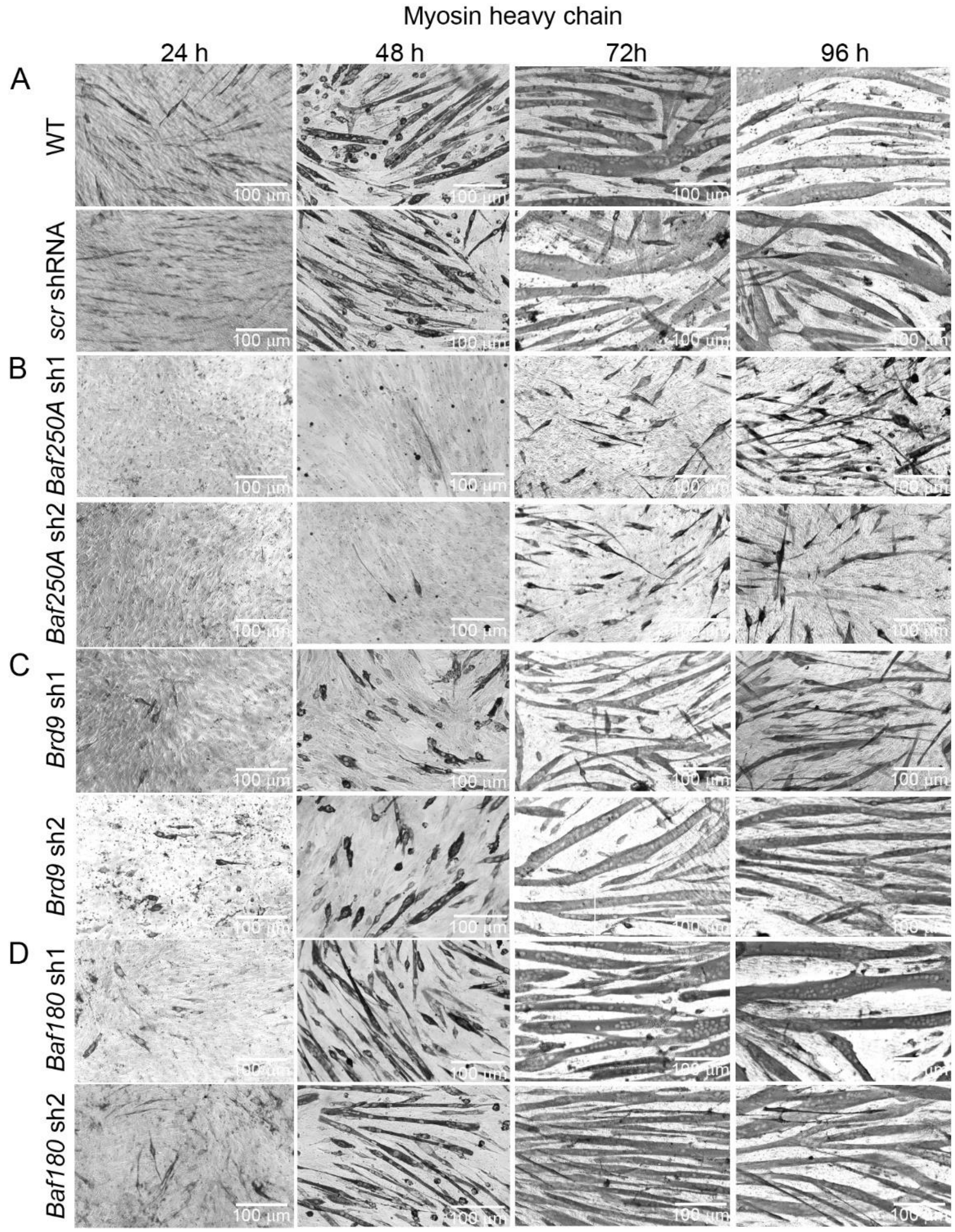


**Supp. Fig. 1. *Baf250A* knockdown inhibited the differentiation of C2C12 cells.** Representative light micrographs of differentiating **(A)** wild type (WT) C2C12 myoblasts and cells transduced with *scr* shRNA **(B)** transduced with *Baf250A* shRNAs, **(C)** transduced with *Brd9* shRNAs, or **(D)** transduced with *Baf180* shRNAs. Cells at 24, 48, 72 and 96 h of differentiation were immunostained for myosin heavy chain. Bars = 100 µm.

### Supplementary Figure 2

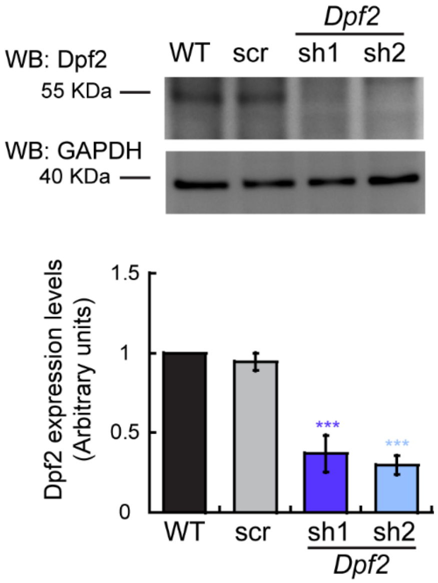


**Supp. Fig. 2. Expression levels of Dpf2 in differentiating wildtype (WT) C2C12 myoblasts.** Representative immunoblots (top) and quantification of Dpf2 levels in myoblasts differentiating for 48 h. Immunoblots against GAPDH were used as loading controls. Samples were compared to the wild type sample, the value of which was set at 1.0. Data are the mean ± SE for three independent experiments. ***P < 0.001.

### Supplementary Figure 3

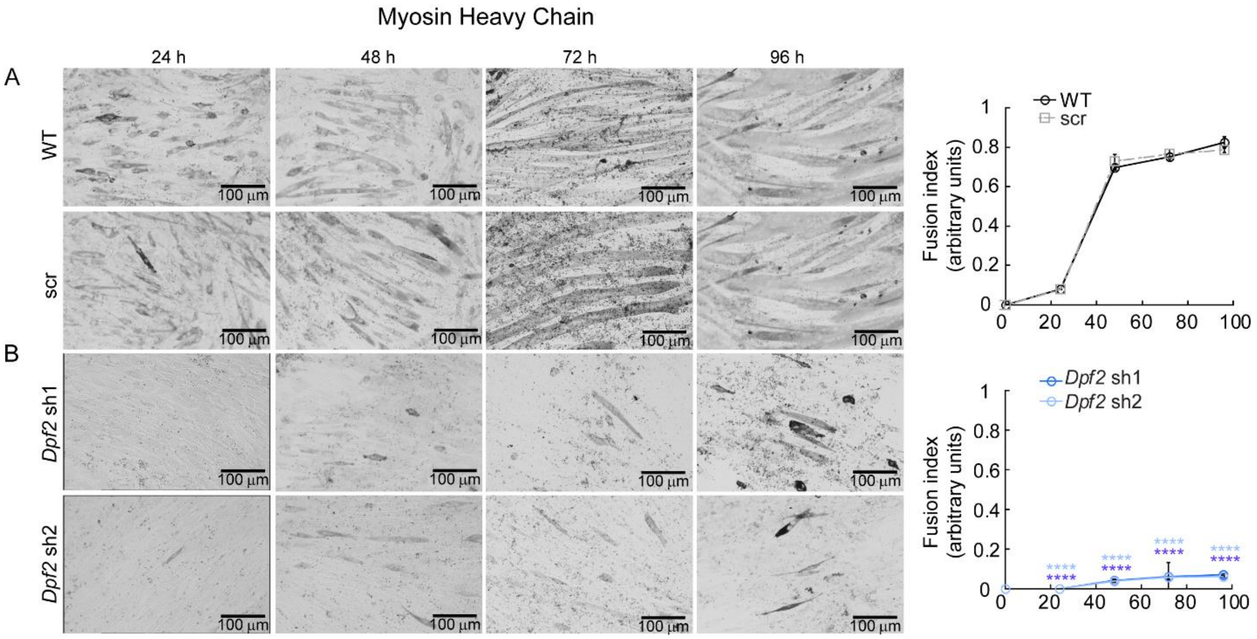


**Supp. Fig. 3.** ***Dpf2* knockdown inhibited the differentiation of C2C12 cells.** Representative light micrographs and fusion index of differentiating **(A)** wild type (WT) C2C12 myoblasts and cells transduced with *scr* shRNA **(B)** transduced with *Dpf2* shRNAs. Cells at 24, 48, 72 and 96 h of differentiation were immunostained for myosin heavy chain. Bars = 100 µm. **** $P < 0.0001$.

**Supplemental Table 1. Sequences of the shRNAs used in this study**

| Gene name | Sequence | Catalog number (Sigma) |
|---|---|---|
| *Baf250* sh1 | CCGGCTTTATAGTATGGCGAGTTAACTCGAGTTAACTCGCCATACTATAAAGTTTTTG | TRCN0000238304 |
| *Baf250* sh2 | CCGGCCTAGGCAGCCTAACTATAATCTCGAGATTATAGTTAGGCTGCCTAGGTTTTTG | TRCN0000238306 |
| *Brd9* sh1 | CCGGTGGACTTTGGCACGATGAAAGCTCGAGCTTTCATCGTGCCAAAGTCCATTTTTG | TRCN0000225737 |
| *Brd9* sh2 | CCGGCACCGAATGGTGTCCAATAAGCTCGAGCTTATTGGACACCATTCGGTGTTTTTG | TRCN0000225739 |
| *Baf180* sh1 | CCGGTGTGAAGTTGGTCCTAGTTTACTCGAGTAAACTAGGACCAACTTCACATTTTTG | TRCN0000304680 |
| *Baf180* sh2 | CCGGGTGCAATATCCAGACTATTATCTCGAGATAATAGTCTGGATATTGCACTTTTTG | TRCN0000304681 |
| *Dpf2* sh1 | CCTGGTGATTACAGGGTCAAA | TRCN0000084343 |
| *Dpf2* sh2 | GCCTAACAACTACTGTGACTT | TRCN0000084345 |
| *scr* shRNA pLKO.1-puro non-target shRNA control plasmid DNA | CCGGCAACAAGATGAAGAGCACCAACTCGAGTTGGTGCTCTTCATCTTGTTGTTTTT | MFCD07785395 SHC002 |

**Supplemental Table 2. List of primers used in this study**

| Primer name | 5' sequence | 3' sequence | Use | Reference |
|---|---|---|---|---|
| *q-Myogenin* | CAAGTGTGCACATCTGTTCTAGTCTCT | GTATCATCAGCACAGGAGACCTTGGT | Gene expression | Hernandez-Hernandez *et al.*, 2013[1] |
| *q-Ckm* | CTGTCCGTGGAAGCTCTCAACAGC | TTTTGTTGTCGTTGTGCCAGATGCC | Gene expression | Hernandez-Hernandez *et al.*, 2013[1] |
| *q- MyHCIIb* | TCAATGAGATGGAGATCCAGCTGAAC | GTCCAGGTGCAGCTGTGTGTCCTTC | Gene expression | Hernandez-Hernandez *et al.*, 2013[1] |
| *q-Cav3* | TCAATGAGGACATTGTGAAGGTAGA | CAGTGTAGACAACAGGCGGT | Gene expression | Witwicka *et al.*, 2019[2] |
| *q-Eef1A1* | GGCTTCACTGCTCAGGTGATTATC | ACACATGGGCTTGCCAGGGAC | Gene expression | Hernandez-Hernandez *et al.*, 2013[1] |
| *Myogenin* promoter | ACGCCAACTGCTGGGTGCCA | GAATCACATGTAATCCACTGGA | ChIP qPCR | Hernandez-Hernandez *et al.*, 2013[1] |
| *Ckm* enhancer | GACACCCGAGATGCCTGGTT | GATCCACCAGGGACAGGGTT | ChIP qPCR | Hernandez-Hernandez *et al.*, 2013[1] |
| *MyHCIIb* promoter | CACCCAAGCCGGGAGAAACAGCC | GAGGAAGGACAGGACAGAGGCACC | ChIP qPCR | Hernandez-Hernandez *et al.*, 2013[1] |
| *Cav3* promoter | CCTAGGTGTCTCAGTCCAGTTA | CTGCCACGTAGATCTTGGAAAT | ChIP qPCR | Witwicka *et al.*, 2019[2] |
| *IgH* enhancer | GCCGATCAGAACCAGAACACC | TGGTGGGGCTGGACAGAGTGTTTC | ChIP qPCR | Hernandez-Hernandez *et al.*, 2013[1] |

**Supplemental Table 3. Pearson coefficients for the two replicate samples for each KD RNA-Seq dataset**

| Sample | Pearson coefficient |
|---|---|
| *scr* shRNA differentiation | 0.96 |
| *Baf250A* shRNA differentiation | 0.98 |
| *Brd9* shRNA differentiation | 0.965 |
| *Baf180* shRNA differentiation | 0.975 |
